# Varenicline Prevents SARS-CoV-2 Infection In Vitro and in Rhesus Macaques

**DOI:** 10.1101/2021.06.29.450426

**Authors:** Jeffrey Nau, Priya Luthra, Kathleen Lanzer, Frank Szaba, Tres Cookenham, Eric Carlson

## Abstract

**Background:** SARS-CoV-2 infections have resulted in a global pandemic, but an antiviral therapy for this novel strain of coronavirus does not currently exist. The objective of our study was to investigate the antiviral potential of the nicotinic acetylcholine receptor (nACHR) agonist varenicline tartrate against SARS-CoV-2.

**Methods:** We assessed antiviral activity using *in vitro* human cell assays and we assessed *in vivo* efficacy in a rhesus macaque model.

**Results:** *In vitro* studies found that varenicline tartrate, over a range of concentrations, reduced the infectivity of SARS-CoV-2 wildtype, alpha, and beta variants in Calu-3 cells and Caco-2 cells, with maintenance of cell viability. *In vivo* studies found that varenicline tartrate, administered as a nasal spray to rhesus macaques, reduced SARS-CoV-2 wildtype viral load and inhibited viral replication in the nasal mucosa and upper airway.

**Conclusion:** Although the study reported here was exploratory, we have confirmed that the nAChR agonist varenicline has the potential to interact with and inhibit SARS-CoV-2 infection and replication.

## INTRODUCTION

The severe acute respiratory syndrome coronavirus-2 (SARS-CoV-2) is a novel strain of coronavirus that was first identified as an infectious agent in humans in 2019.^1^ The disease caused by the SARS-CoV-2 viral outbreak was termed coronavirus disease 2019 (COVID-19); the major clinical manifestations include cough, shortness of breath or difficulty breathing, fever, chills, muscle pain, headache, sore throat and a new loss of taste or smell. This viral outbreak was recognized as a pandemic threat by the World Health Organization and began to rapidly overload health care systems, causing substantial morbidity and mortality worldwide.^2,3^ Recent evidence reports from 600,000-900,000 deaths in the United States, and from 3.9-7.1 million deaths worldwide.^4,5^ Importantly, an effective preventative antiviral treatment for SARS-CoV-2 has not yet emerged.^6^ Compounding the urgency for an effective treatment is the emergence of mutations (with changes in the spike protein) with increased rates of transmission and mortality.^7,8^ For example, the beta variant (first identified in South Africa) has 50% increased transmission and reduced neutralization by post-vaccination antibodies compared with wildtype virus,^9^ and the alpha variant (first identified in the United Kingdom) is more transmissable than the wildtype and has a 62.8% increase in mortality after 28 days.^10^

Many studies have shown one route that SARS-CoV-2 uses to enter the host cell is via binding to angiotensin-converting enzyme-2 (ACE2) and subsequent fusion with the cell membrane.^11-13^ Additional studies have shown that SARS-CoV-2 enters host cells mainly through ACE2-positive and transmembrane serine protease 2-positive nasal epithelial cells, which are specific ciliated and mucous secreting cells of the nasal mucosa.^14,15^ Therefore, the nasal cavity potentially represents the most susceptible mucosal surface for infection within the respiratory tree, although the ocular surface is also susceptible to infection.^16^ Concurrently, based on early observations of low smoking prevalence among hospitalized COVID-19 patients in China and the USA, a role for the nicotinic cholinergic system in SARS-CoV-2 infection was postulated.^17,18^ However, more recent studies suggest the relationship between smoking and SARS-CoV-2 is complex, and that smoking may indeed be a risk factor for severe disease,^19^ but the role of nicotine and the nicotinic cholinergic system remains pertinent, though unclear.

Farsalinos et al. have identified a “toxin-like” amino acid in the receptor binding domain of the spike glycoprotein of SARS-CoV-2, that is homologous to a sequence found in snake venom known to interact with nicotinic acetylcholine receptors (nAChRs).^20,21^ Further, binding simulation studies have shown that SARS-CoV-2 has the ability to directly bind to nAChRs via its spike protein.^22^ Binding to the nAChRs may explain the immune-mediated flares or myasthenia gravis-like findings in patients with autoimmune disease infected with SARS-CoV-2, or those vaccinated with mRNA/DNA based vaccines.^23,24^ Receptor binding raises the possibility that cholinergic agonists may have the ability to block nAChR interaction with SARS-CoV-2, as well as other coronaviruses that share similar amino acid sequences in their receptor binding domain. To complicate this potential role, the nicotine cholinergic system may also interact with the ACE2 system.^19^ Overall, it is possible that nAChR agonists may have therapeutic value in SARS-CoV-2 infected patients, making them worthy of study.

Varenicline tartrate is a highly selective small-molecule nAChR agonist with full agonist activity at the α7 receptor and partial agonist activity at the α3β4, α3α5β4, α4β2, and α6β2 receptors.^25-28^ Varenicline tartrate is FDA approved and indicated for use as an aid to smoking cessation treatment in oral tablet form^29^ and is currently under investigation as a preservative-free aqueous nasal spray formulation (OC-01) for dry eye disease. In this study, we hypothesized that varenicline may have antiviral activity against SARS-CoV-2 by inhibiting viral host cell entry and thereby disrupting the replication cycle. Of note, varenicline in particular has been shown, using computational strategies, to bind to the spike glycoprotein of SARS-CoV-2 at the hinge site with high affinity.^30^ By administering as a topical formulation, OC-01 (varenicline) nasal spray of offers a unique opportunity to target SARS-CoV-2 at the primary site of infection, binding, amplification, and transmission. Topical administration of varenicline as an aqueous nasal spray allows for the delivery of a high localized concentration to the mucosa that is unlikely to be achieved with systemic delivery. The aim of this initial exploratory study was to investigate the antiviral potential of varenicline against SARS-CoV-2 using *in vitro* cell assays and an *in vivo* rhesus macaque model. Prior studies with SARS-CoV-2 wildtype using rhesus macaque model have found that infection results in mild to moderate clinical signs and high titers of virus in the upper and lower respiratory tract with multifocal lung inflammation.^31^

## METHODS

### *In vitro* antiviral activity in Calu-3 and Caco-2 cells

#### Reagents

Antiviral activity assays were conducted in Calu-3 and Caco-2 cells. Calu-3 cells (ATCC ATB-55) are a line of epithelial cells derived from a lung adenocarcinoma and Caco-2 cells (ATCC HTB-37) are a line of epithelial cells from a colorectal adenocarcinoma: both are anticipated to express both ACE2 and nAChR receptors.^32,33^ SARS-CoV-2 wild-type clinical isolate, USA WA1/2020, SARS-CoV-2 alpha (B.1.1.7 UK), and SARS-CoV-2 beta (B.1.351 South Africa) clinical isolates were obtained from BEI Resources. All viruses were passaged in Vero E6 cells to create the virus working stock. Varenicline tartrate was purchased from Tocris Biosciences (Minneapolis, MN). Varenicline tartrate was reconstituted in DMSO to yield stock solutions of 20 mM. Stocks were stored at -20 °C, thawed immediately before the experiment and residual compound was discarded. All the work with SARS-CoV-2 was performed in a BSL-3 laboratory at the Trudeau Institute following institutional approved biosafety protocols.

#### Experimental procedure

A 3-fold dilution series of varenicline tartrate was prepared starting at 100 μM. Cells were pre-treated with varying concentrations of varenicline tartrate for 1 hour. Separately, SARS-CoV-2 was diluted to generate a multiplicity of infection (MOI) of 3 (3 virions per cell) and was pre-treated with various concentrations of varenicline tartrate (control = DMSO) for 1 hour at 37° C and under 5% CO_2_. Following the pre-treatment period, cells were exposed to the pre-treated SARS-CoV-2/varenicline tartrate for 2 hours and the plates were shaken every 15 minutes. Following this 2 hour incubation, the culture medium was removed and the cells were washed twice with PBS. Fresh culture medium containing various concentrations of varenicline tartrate (DMSO served as a negative) was then added to the cells. Cell culture supernatants were harvested at 24 hours post SARS-CoV-2/varenicline tartrate exposure and subjected to plaque titration. For plaque titration, confluent Vero-E6 cells were inoculated with 5-fold serial dilutions of cell culture supernatant and incubated for 1 hour at 37°C and 5% CO_2_. Following this incubation period, culture medium containing 2.4% (final concentration 0.6%) microcrystalline cellulose was added to the cultures. Plaques were counted 72 hours post-infection and titers determined as plaque forming units per ml (pfu/ml). The percent infection of cells was normalized against virus only control samples. Cell viability was determined at 72 hours using the CellTiter-Glo Luminescent Cell Viability Assay kit (Promega) and luminescence was recorded using a POLARstar OPTIMA plate reader (BMG Labtech). The percent cell viability was calculated by normalizing against DMSO only controls.

#### Statistical analyses

All statistical analyses were performed using GraphPad Prism (version 9, GraphPad Software, Inc.). For the calculation of 50% inhibitory concentration (IC_50_) and 50% cytotoxic concentration (CC_50_) of the compound, which indicate the inhibitor concentration leading to 50% reduction of infection or cell viability respectively, non-linear fit regression models were used.

### In vivo SARS-CoV-2 challenge study in Rhesus Macaques

Studies were conducted under the BIOQUAL Inc. Institute Institutional Animal Care and Use Committee (IACUC)-approved protocol 20-069P, which is in compliance with the Animal Welfare Act and other federal statutes and regulations relating to animals and experiments involving animals. BIOQUAL Inc. is accredited by the Association for Assessment and Accreditation of Laboratory Animal Care International and adheres to principles stated in the Guide for the Care and Use of Laboratory Animals, National Research Council.

#### Experimental procedure

To evaluate the ability of varenicline OC-01 nasal spray to prevent nasal infection and replication in rhesus macaques from SARS-CoV-2, we assessed the impact of varenicline administration prior to, and following virus challenge (Figure 1). Each animal (N = 2) was treated with 100 µL of 0.6 mg/ml varenicline OC-01 nasal spray into each nostril 4 times daily for 5 days (Days 1-5). On Day 1, the first and second OC-01 doses were administered, then animals were challenged with 1×10^5^ pfu of SARS-CoV-2 WA/1/2020 isolate through intranasal and intratracheal routes. Prior to inoculation, the same lot of isolate (Lot# 70040665 NR-53872) was tested for infectivity in a titration experiment in the rhesus macaque (N = 3) control animals. Then the third and forth OC-01 doses were administered. On Days 2 to 5, 4 doses of OC-01 were given, and on Day 6, 2 doses of OC-01 were given, and then animals were euthanized (with an intravenous overdose of sodium pentobarbital) for necropsy and tissue collection. Based on the relatively transient nature of the SARS-CoV-2 infection in rhesus macaques, the in-life portion of the study was limited to 6 days.

**Figure 1.**
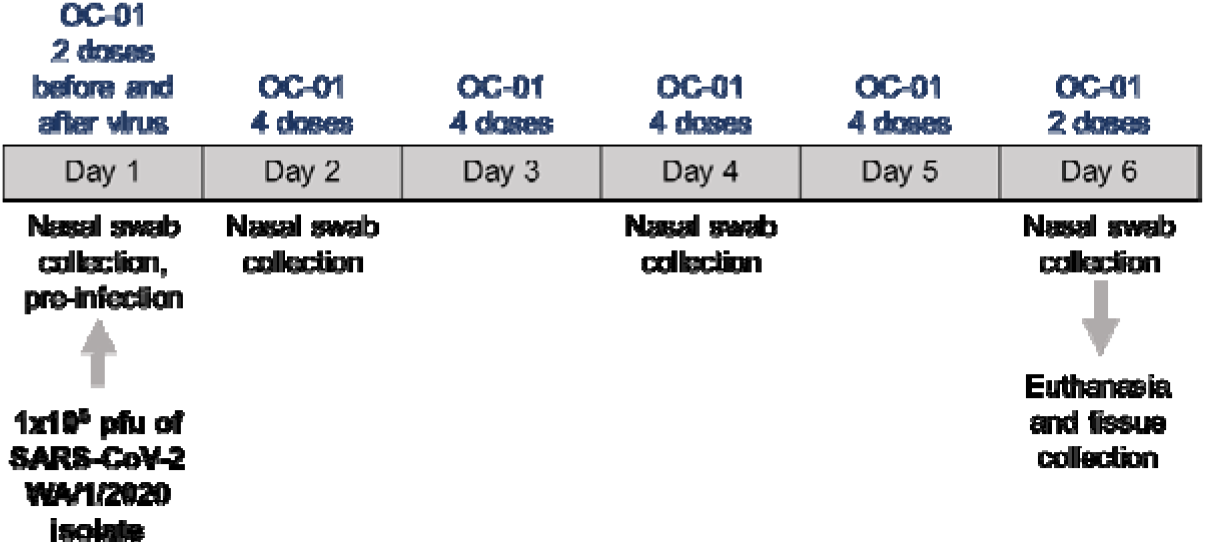
Study Design for SARS-CoV-2 Challenge in Rhesus Macaques. Each animal (N = 2) was treated with varenicline OC-01 nasal spray into each nostril 4 times daily for 5 days. On Day 1, the first and second OC-01 doses were administered, then animals were challenged with 1×10^5^ pfu of SARS-CoV-2 WA/1/2020 isolate, then the third and forth OC-01 doses were administered. On Days 2 to 5, 4 doses of OC-01 were given, and on Day 6, 2 doses of OC-01 were given, and then animals were euthanased. Nasal swabs were collected for viral load analysis by qRT-PCR. Control animals (N = 3) did not received OC-01 treatment.

Animals were given enrichment, including commercial toys and food supplements. All animals were monitored for clinical signs and body weight was assessed. To determine viral load in upper airway, nasopharyngeal swabs were collected on Days 2, 3, 5, and 6 for control animals and Days 2, 4, and 6 for OC-01-treated animals. Collected swabs were stored at -80°C until viral load analysis.

#### Determination of viral load

##### Quantitative RT-PCR for SARS-CoV-2 genomic (g) RNA

Samples were assayed for virus concentration by SARS-CoV-2-specific qRT-PCR, which estimated the number of RNA copies in the collected tissue samples. Viral RNA was isolated from collected tissue samples using the Qiagen MinElute virus spin kit and a control for the amplification reaction was isolated from the SARS-CoV-2 stock. qRT-PCR was performed with Applied Biosystems 7500 Sequence detector and amplified using the following program: 48°C for 30 minutes, 95°C for 10 minutes followed by 40 cycles of 95°C for 15 seconds, and 1 minute at 55°C using primers (2019-nCoV_N1-F :5’-GACCCCAAAATCAGCGAAAT-3’; 2019-nCoV_N1-R: 5’-TCTGGTTACTGCCAG TTGAATCTG-3’) and a probe (2019-nCoV_N1-P: 5’-FAM-ACCCCGCATTACGTTTGGTGGACC-BHQ1-3’) specifically designed to amplify and bind to a conserved region of nucleocapsid gene of coronavirus. The signal was compared to a known standard curve and calculated to give copies per mL (in a practical range of 50 to 5 × 10^8^ RNA copies per mL for nasal washes).

##### Quantitative RT-PCR for SARS-CoV-2 subgenomic (sg) RNA

Prior to PCR, samples collected from animals were reverse-transcribed using Superscript III VILO (Invitrogen). A Taqman custom gene expression assay (ThermoFisher Scientific) was designed using the sequences targeting the E gene sgRNA20 (SG-F: CGATCTTGTAGATCTGTTCCTCAAACGAAC; SG-R: ATATTGCAGCAGTACGCACACACA; probe: FAM-ACACTAGCCATCCTTACTGCGCTTCG-BHQ) and PCR reactions were carried out on a QuantStudio 6 and 7 Flex Real-Time PCR System (Applied Biosystems). To generate a standard curve, the SARS-CoV-2E gene sgRNA was cloned into a pcDNA3.1 expression plasmid; this insert was transcribed using an AmpliCap-Max T7 High Yield MessageMaker Kit (Cellscript) to obtain RNA for standards. Standard curves were used to calculate sgRNA in copies per swab (in a practical range of 50 to 5 × 10^7^ RNA copies per mL for nasal washes).

## RESULTS

### Varenicline tartrate inhibits SARS-CoV-2 *in vitro*

We assessed the antiviral activity of varenicline tartrate against SARS-CoV-2 in Calu-3 and Caco-2 cells. Varenicline tartrate reduced SARS-CoV-2 viral titers over a range of concentrations, with an IC_50_ of 0.3 μM and 0.5 μM in Calu-3 (Figure 2A) and Caco-2 cells (Figure 2B), respectively. In addition, varenicline tartrate reduced SARS-CoV-2 alpha variant viral titers with an IC_50_ of 0.13µM in Calu-3 cells (Figure 2C), and SARS-CoV-2-beta variant viral titers with an IC_50_ of 4µM in Calu-3 cells (Figure 2D). Importantly, these *in vitro* conditions were not toxic and cell viability was maintained.

**Figure 2.**
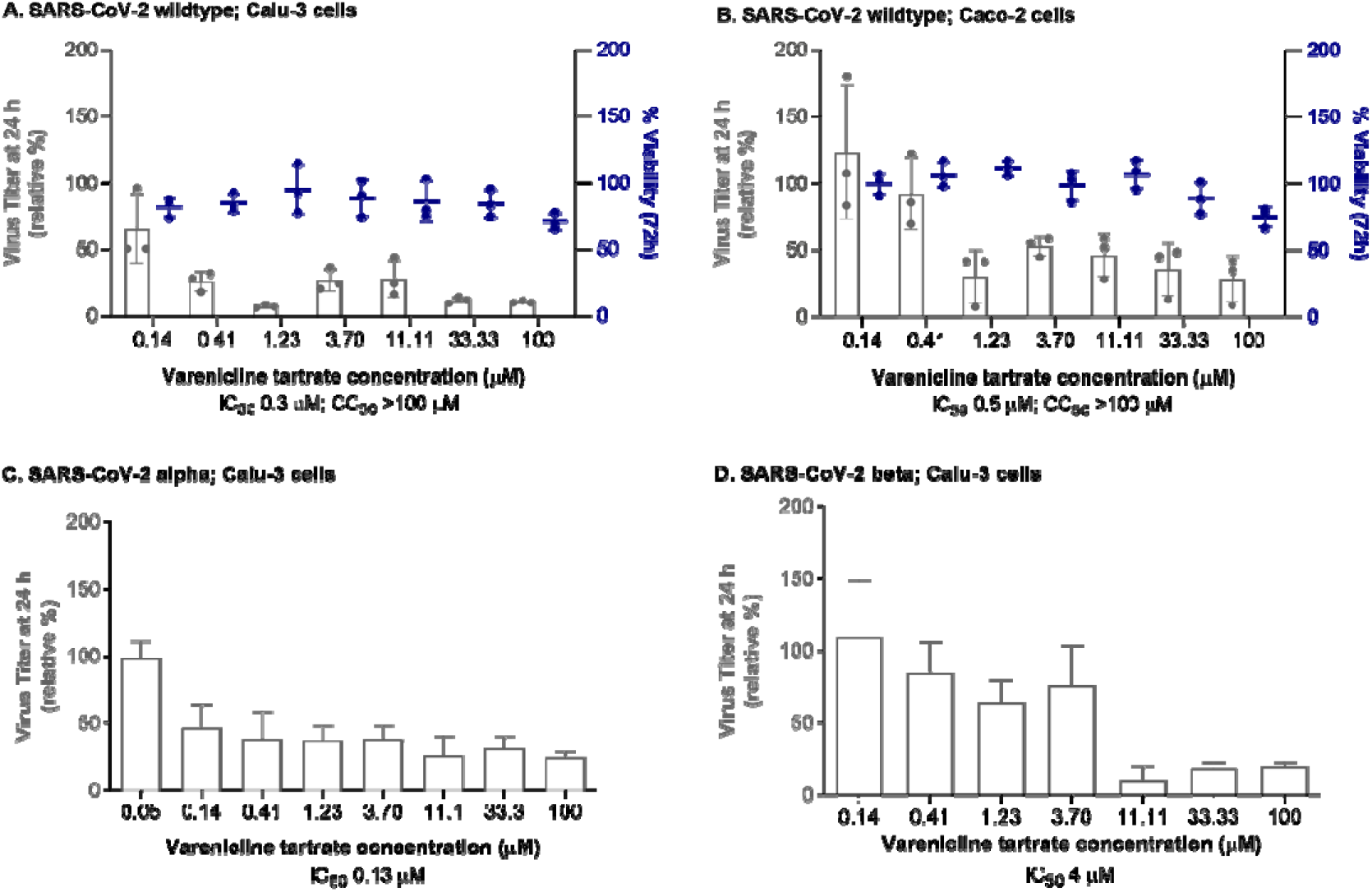
Antiviral activity of varenicline tartrate in Calu-3 and Caco-2 cells. The target cells, Calu-3 and Caco-2, as well as SARS-CoV-2 (wildtype, alpha, and beta variants) were preincubated with different concentrations of varenicline tartrate for 1h at 37°C. The cells were infected with pre-treated virus at an MOI of 3 and antiviral activity was assessed by using the cell supernatants 24h post-infection. Error bars represent mean and standard deviation of three independent replicates: the experiment was performed as two independent replicates. CC_50_ 50% cytotoxic concentration; IC_50,_ 50% inhibitory concentration.

### Varenicline tartrate inhibits SARS-CoV-2-wt *in vivo*

We assessed the antiviral activity of varenicline against SARS-CoV-2 in rhesus macaques. During the study, there were only minor changes in body weight and rectal temperatures, and no changes in respiratory rate, respiratory effort, cough, and fecal consistency (data not shown). No adverse events were detected.

Nasopharyngeal swabs were assessed for SARS-CoV-2 by qRT-PCR for gRNA, which indicates the presence of virus, and sgRNA (sgRNA), which indicates the presence of actively replicating virus. In control animals, following SARS-CoV-2 infection, copies of viral RNA peaked approximately 2 days post-challenge (Figure 3A and B). Varenicline OC-01 nasal spray prevented significant SARS-CoV-2 infection in the nasal mucosa of the rhesus macaques (Figure 3C and D). Following treatment with OC-01, SARS-CoV-2 gRNA levels were decreased approximately 100-fold (Figure 3C), and sgRNA levels were decreased approximately 200-fold compared with controls. sgRNA levels were no longer detectable at 4 days and 6 days post-challenge (Figure 3D).

**Figure 3.**
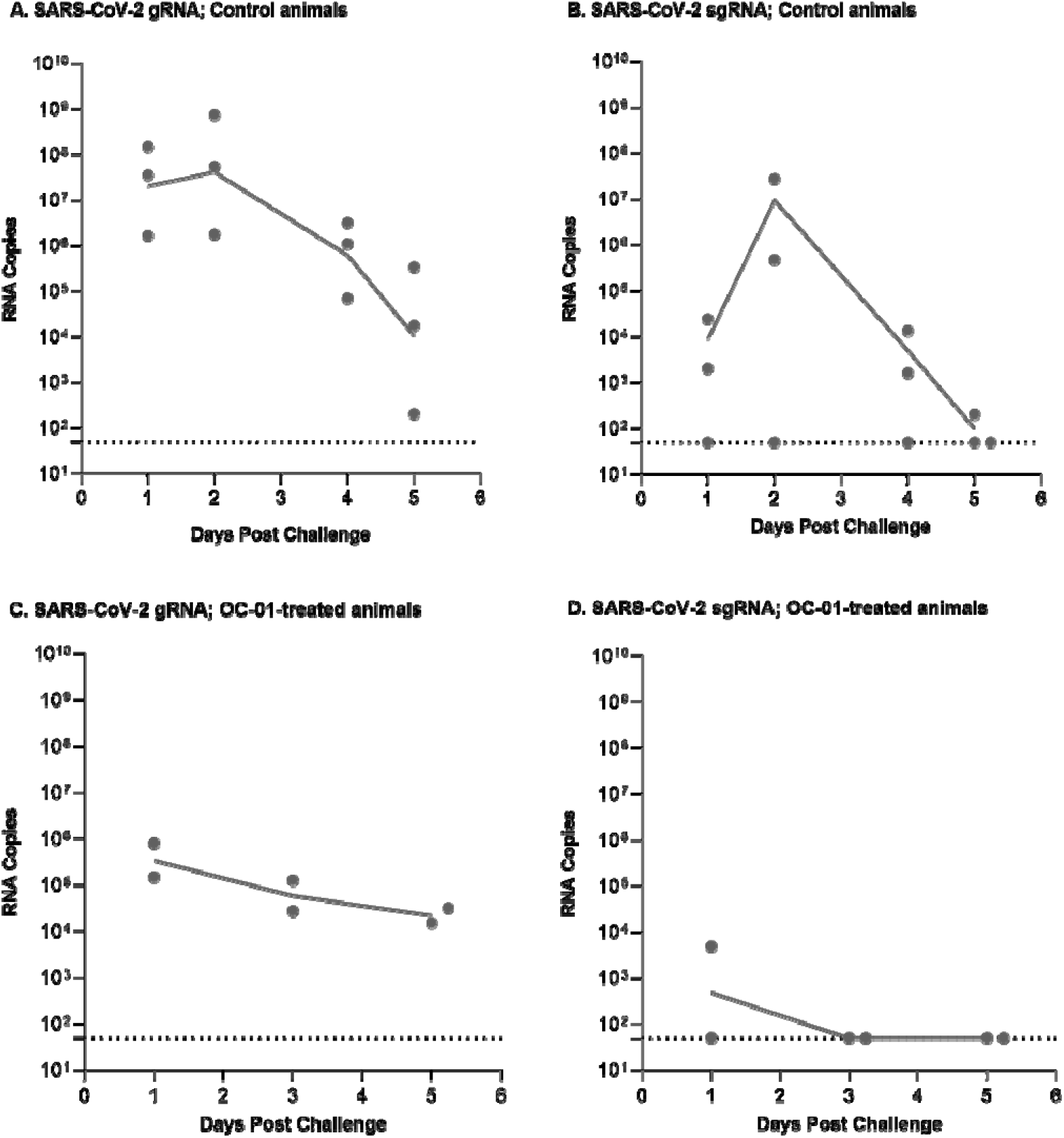
Effect of varenicline nasal spray on SARS-CoV-2 gRNA and sgRNA in rhesus macaques. SARS-CoV-2 gRNA and sgRNA were measured by qRT-PCR in nasal swab samples taken at Day 2, Day 3, Day 5, and Day 6 of the study (equivalent to post-challenge days 1, 2, 4, and 5) for the control animals and Day 2, Day 4, and Day 6 of the study (equivalent to post-challenge days 1, 3, and 5) for the OC-01-treated animals. Animals were challenged with SARS-CoV-2 on Day 1 of the study, which is equivalent to 0 days post challenge in the figure. Dotted lines represent lowest level of detection for the assay.

## DISCUSSION

To our knowledge these are the first *in vitro* and *in vivo* studies to show that a nAChR agonist has antiviral activity against SARS-CoV-2. Our initial *in vitro* cell assays demonstrated that varenicline reduced viral titers over a range of doses, against wildtype, alpha, and beta variants, without a negative impact on cell viability. Our initial *in vivo* studies indicated that varenicline administered as an aqueous nasal spray (approximately 1mM at each administration based on airway surface liquid calculations) prevented SARS-CoV-2 infection and replication in the nasal cavity of rhesus macques less than 24 hours after virus challenge. The results suggest a sound rationale for the use of OC-01 (varenicline) nasal spray as a therapeutic for pre-exposure/post-exposure prophylaxis to prevent infection, to decrease viral load, and/or to lessen severity and transmission of SARS-CoV-2. Additionally, *in vitro* results suggest that the mechanism for viral inhibition is conserved with the SARS-CoV-2 alpha and SARS-CoV-2 beta variants. Moreover, the nasal spray formulation of varenicline enables administration of a therapeutic dose directly to the nasal mucosa, which represents the most likely route of virus entry, replication, and infection for SARS-CoV-2.^12,22,34-36^ In addition, there is a significant population of nAChRs on the olfactory bulb within the nasal mucosa;^37^ this may help to understand the chemo-sensitive disorders seen with SARS-CoV-2 infection, such as loss or decline of taste and smell (ageusia and anosmia, respectively), which have been reported as unique clinical features of COVID-19.^38,39^

Our results are consistent with data from *in silico* studies that suggested varenicline binds directly to the receptor binding domain of the spike glycoprotein of SARS-CoV-2 at the hinge site with high affinity. This binding may prevent a change in the spike protein to the “up-conformation” and inhibit subsequent binding by the ACE2 and/or nAChR binding site,^30^ thus potentially preventing host cell infection. In addition, the spike glycoprotein of SARS-CoV-2 may also directly bind with nAChRs, and binding studies have suggested that the Y674-R685 region of the protein adopts particular confirmations when binding to the α4β2 and α7 nAChR subtypes,^22^ which also have the same high affinity for varenicline.^25^ Therefore, unlike anti-SARS-CoV-2 vaccines and antibody therapeutics, the antiviral activity of varenicline is likely to be conserved for different SARS-CoV-2 variants as varenicline’s affinity for nAChRs or the Y674-R685 region of the spike protein is completely independent of virus mutations.^25-28,30^

Although the studies reported here were exploratory, we have confirmed that the nAChR agonist varenicline, the active ingredient of OC-01 nasal spray, has the potential to interact with and inhibit SARS-CoV-2 infection and replication. Further, the nicotinic cholinergic system has been postulated to be involved in the pathophysiology of severe COVID-19 due to the immune dysregulation and cytokine storm, as the cholinergic anti-inflammatory pathway may be an important regulator of the inflammatory response.^17,40^ Given the *in vitro* and *in vivo* effectiveness seen in the studies, varenicline nasal spray warrants further investigation as an antiviral agent for pre-exposure/post-exposure prophylaxis, and/or prevention of transmission of SARS-CoV-2 wildtype and variants.

## Role of contributors

EC and JN participated in the study design, interpretation of study results, and drafting of the manuscript. All authors participated in the critical revision and approval of the final version of the manuscript. PL, KL, FS and TC were involved in the *in vitro* study execution, data collection, and data analysis.

